# Scalable Decoupling Contact and Configuration Spaces in a Quantum-Enabled Protein Folding Workflow

**DOI:** 10.64898/2026.10.09.758014

**Authors:** Akshay Uttarkar, Vidya Niranjan

**Author notes:** Corresponding author: Vidya Niranjan.

## Abstract

Near-term quantum devices remain constrained by limited qubit counts, noise, and shallow circuit depth, restricting their direct application to peptide folding. This work addresses this limitation through a hybrid quantum–classical framework that separates global contact-space exploration from local configuration-space refinement.

The pipeline was evaluated on the p53 transactivation peptide. The peptide was encoded using discrete torsional states and Miyazawa–Jernigan contact potentials. Global conformational search was performed using simulated annealing, followed by fragment-level CVaR-based variational quantum eigensolver refinement. Overlapping fragments were stitched and reconstructed into three-dimensional backbone coordinates for structural comparison.

Three independent simulated annealing restarts produced negative-energy conformers. The workflow identified a best global conformer at E = −3.70 MJ using simulated annealing. Fragment-level CVaR-VQE refinement generated 30 candidate conformers across three overlapping fragments, the best assembled conformer reaching E = −2.58 MJ after DP stitching and post-stitch refinement. In this trial, the classical SA solution outperformed the VQE-assembled structure and was selected as the final output. The final 65-atom model was successfully reconstructed, but full-backbone alignment against 1YCR yielded an RMSD of 4.65 Å, improving to 2.90 Å after trimmed alignment.

Our work demonstrates a practical quantum-enabled peptide folding workflow for short, flexible peptides, but the present structural accuracy is insufficient for claims of native fold recovery. Its main value is as an auditable framework for testing NISQ-compatible folding strategies.

## 1. Introduction

The prediction of a protein’s three-dimensional native structure from its primary linear sequence of amino acids is a central unsolved problem in computational biology. All structural prediction algorithms, classical and quantum alike, rest on the thermodynamic hypothesis (Anfinsen’s dogma): the native, biologically active conformation corresponds to the global free-energy minimum under physiological conditions. [1] Protein folding therefore reduces to a high-dimensional global optimization problem: locating the lowest-energy atomic arrangement within a rugged energy landscape.

The geometric vastness of a protein’s conformational space poses an extreme barrier to computational simulation, as Levinthal famously noted in 1969 [2]. Levinthal observed that the backbone of a protein possesses specific dihedral degrees of freedom (primarily the phi and psi angles between adjacent residues). For a modest protein of 100 amino acids, the number of possible configurations is estimated conservatively at 10^300^ or greater [3].

Finding the exact lowest free-energy conformation of a protein is an NP-hard combinatorial optimisation problem [1]. While it is trivial to verify the energy of a proposed structure once given, searching the full conformational space scales exponentially with sequence length [4]. Classical molecular dynamics (MD) simulations attempt to resolve this by integrating Newton’s equations of motion over discrete femtosecond time steps, but even modern supercomputers require hours to simulate nanoseconds of molecular motion [3]. Since biologically relevant folding occurs on millisecond timescales, classical MD cannot bridge this gap [5]. Heuristic methods such as Monte Carlo and simulated annealing help but still get trapped in local minima [5]. This intractability motivates quantum computational approaches, which can in principle evaluate large superpositions of conformational states simultaneously [6].

To render the NP-hard protein folding problem tractable for mapping onto the limited qubit topologies of current generation quantum processors, continuous three-dimensional Cartesian coordinate space must be discretised into coarse-grained models.[5] Historically, the Hydrophobic Polar (HP) lattice model has served as the foundational testing ground for both classical metaheuristics and novel quantum algorithms [5]. In the HP framework, the sequence is classified into two broad binary categories: hydrophobic (H) beads that seek the protein core, and polar (P) beads that interact favourably with the aqueous solvent. The protein is represented as a self-avoiding chain moving across a tetrahedral lattice [7]. The global energy is calculated by tallying a unit negative energy contribution for every topological instance in which two hydrophobic residues occupy nearest neighbour adjacent lattice sites without being directly connected by the covalent backbone [8]. Despite the extreme abstraction, determining the minimal energy configuration of an HP lattice protein retains NP-hard computational complexity, providing a rigorous benchmark for verifying the success rates of quantum processing units [9].

A central hardware constraint of the current era is the limited number of physical qubits and their coherence lifetimes. As the target protein sequence length N increases, the baseline quantum register size requirement of Q = Ncfg + Nint + 1 expands aggressively [10]. Even with severe coarse-graining, simulating a 10-residue peptide on a tetrahedral lattice requires at least 35 qubits, which exceeds what current hardware supports for deep, error-free gate operations [11]. Robert et al. addressed this by combining classical optimisation with variational quantum models, reducing the qubit requirement for the 10-residue Angiotensin peptide from 35 to 22 [11].

To work within current hardware limits, hybrid pipelines combine classical metaheuristics with targeted quantum subroutines [10]. The hybrid Simulated Annealing and VQE (SA-VQE) pipeline divides the problem into two sequential phases. In the first phase, classical Simulated Annealing surveys the full conformational space using a geometric cooling schedule, applying single spin-flip moves and accepting uphill transitions probabilistically. This pass identifies the most stable macroscopic contact topologies, yielding an optimised vector of contact variables 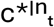 [10]. In the second phase, the optimal contact topology is frozen and passed as static constraints into the VQE Hamiltonian. By projecting the global optimisation problem onto the subspace defined by these fixed variables, 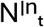 qubits are immediately removed from the necessary quantum register, collapsing the quantum footprint to a highly reduced parameterisation Qreduced = Ncfg + 1, allowing the CVaR VQE algorithm to utilise its coherent entanglement exclusively to resolve the precise backbone stereochemistry that satisfies the static contacts [10]. QuPepFold implements these strategies as a Python-based package for short peptides, with a focus on intrinsically disordered regions (IDRs) relevant to drug targeting [10]. The package takes a user-supplied amino acid sequence, encodes it into discrete configuration variables, evaluates Miyazawa–Jernigan contact energies, and runs multi-start CVaR-VQE optimisation using Nelder–Mead routines within Qiskit Aer. Benchmarking showed the CVaR-optimised architecture reached ground-state conformations roughly 30% faster than a standard expectation-value VQE baseline [12]. On the IonQ Aria trapped-ion processor, the pipeline reproduced absolute ground-state energies with greater than 90% fidelity [12].

These results confirm that hybrid SA-VQE algorithms with CVaR optimisation can operate within current NISQ-era constraints, and establish a computational framework applicable to intrinsically disordered proteins that remain poorly served by classical structure prediction methods [13].

This work addresses this limitation through a hybrid quantum–classical framework that separates global contact-space exploration from local configuration-space refinement. These implementations (Fig 1A) confirm that sophisticated hybrid SA-VQE algorithms, underpinned by rigorous CVaR optimization and precise Miyazawa-Jernigan energetic mapping, can actively overcome current NISQ-era constraints. This methodological foundation opens unprecedented avenues for integrating quantum hardware into advanced drug discovery pipelines, specifically targeting intrinsically disordered proteins that have historically evaded classical computational resolution.

**Figure 1:**
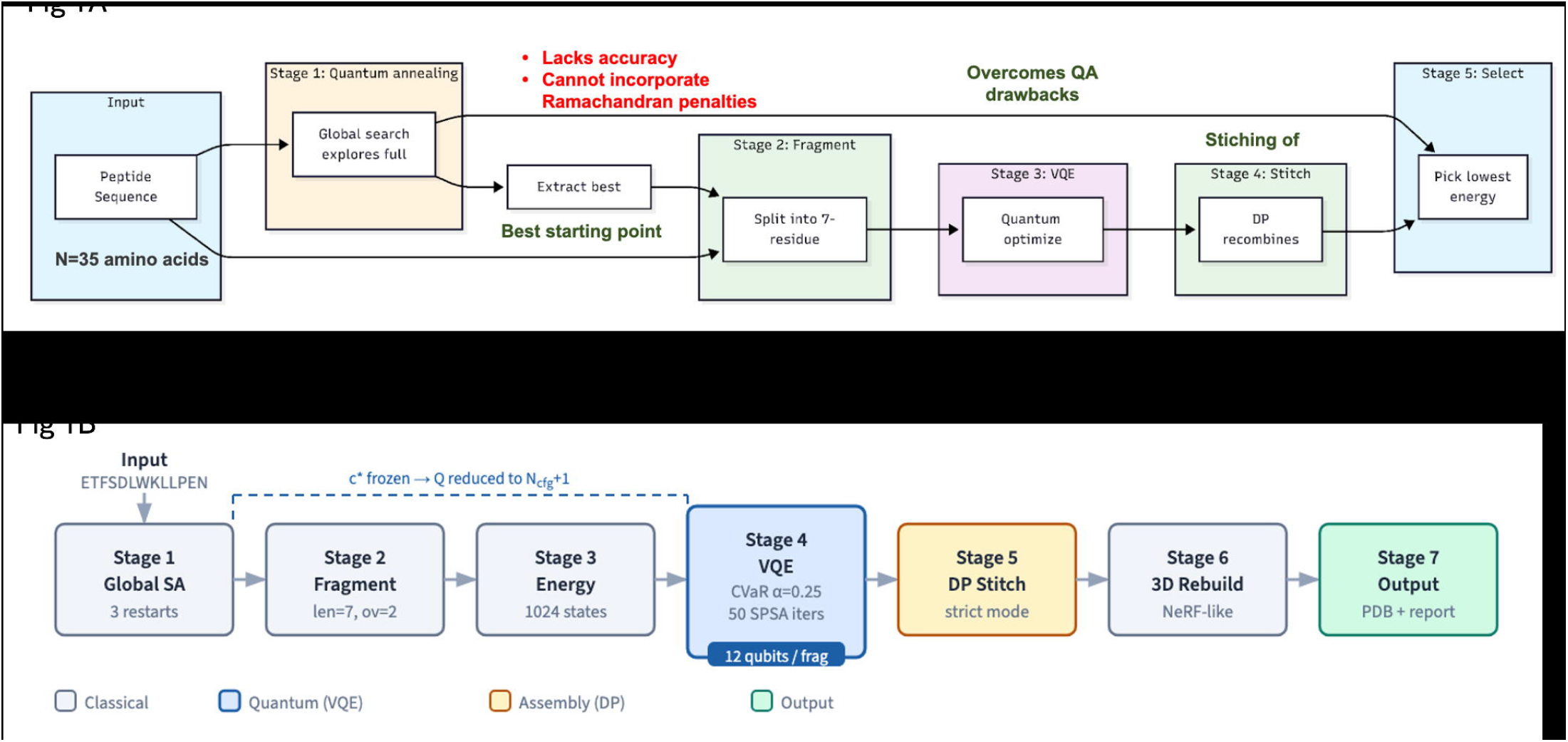
Overview of the QuPepFold hybrid SA-VQE pipeline. A) A schematic of a hybrid quantum–classical framework that separates global contact-space exploration from local configuration-space refinement. B) Schematic flowchart of the 8-stage pipeline: global SA search → fragment generation → energy table precomputation → local VQE refinement (QA-anchored) → dynamic-programming stitching → 3D NeRF-like reconstruction → output. Highlight the qubit-reduction step where the contact topology c* is frozen before VQE.

## 2. Methods

### 2.1 System Modelling and Peptide Lattice Representation

To model the folding behaviour of a peptide sequence of N amino acid residues, we employ a discretised torsional conformation space. Each peptide bond along the backbone possesses dihedral degrees of freedom. We define N−1 discrete turns t_a_ ∈ {0, 1, 2, 3} for a ∈ {0, 1, …, N−2}, mapping each turn state to a specific structural motif:

- *t*_a_ *= 0 ⇒ (*φ_a_, *ψ*_a_*) = (−60°, −45°)* (helix-like conformation)
- *t*_a_ *= 1 ⇒ (*φ_a_, *ψ*_a_*) = (−135°, 135°)* (β strand-like conformation)
- *t*_a_ *= 2 ⇒ (*φ_a_, *ψ*_a_*) = (−75°, 145°)* (polyproline II-like conformation)
- *t*_a_ *= 3 ⇒ (*φ_a_, *ψ*_a_*) = (−60°, 140°)* (extended/coil-like conformation)

Each turn t_a_ is encoded using two binary variables (qubits) 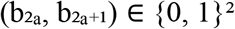 such that:

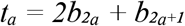

The Kronecker delta functions mapping t_a_ to each discrete state k ∈ {0, 1, 2, 3} are formulated as quadratic polynomials over the binary variables:

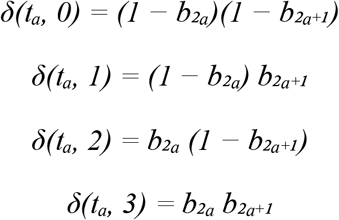

To reduce the active search space and optimise qubit resource requirements, a turn-to-qubit mapping template is applied in which a prefix of length K (e.g., t_0_, …, t_k−1_) is fixed to predetermined ground-state-like folds. The remaining 2(N−1)−2K qubits constitute the active decision variables N_tu_ in the quantum state space.

### 2.2. Peptide Hamiltonian Formulation

The total conformational energy of a given fold is evaluated by a global energy Hamiltonian:

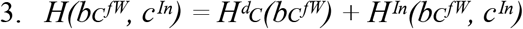

#### 2.2.1 Backbone Conformational Term (H^d^C)

The backbone steric constraint prevents adjacent residues from adopting identical conformational states, modelling local rigidity and chirality:

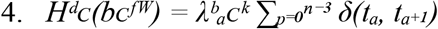

where 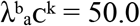 is the backbone penalty coefficient and δ(t_a_, t_a+1_) = Σ_k_ δ(t_a_, k)δ(t_a+1_, k).

#### 2.2.2 Non Local Interaction Term (H^In^)

Non-local residue interactions (hydrophobic/hydrophilic contacts) are represented using binary contact control variables c_ij_ ∈ {0, 1} for residue pairs in the candidate set *P*, defined as:

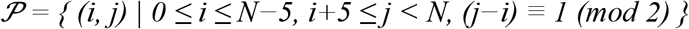

For each pair (i, j) ∈ *P*, c_ij_ is a binary decision variable indicating whether residues i and j form a spatial contact (c_ij_ = 1) or remain separated (c_ij_ = 0).

The interaction Hamiltonian is:

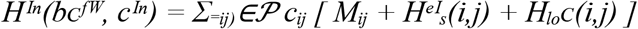

where M_ij_ is the contact energy derived from the Miyazawa–Jernigan (MJ) statistical potential matrix.

#### 2.2.3 Distance and Locality Constraints

To verify the physical feasibility of contacts represented by c_ij_ = 1, we map the relative spatial positions of residues using a 4D representation space. For any residue range [a, b] with a < b, the relative distance vector 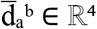 is defined as:

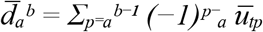

where ū _k_ is the standard basis unit vector in ℝ^4^ corresponding to turn type k. The squared Euclidean norm representing the relative distance between residues a and b is:

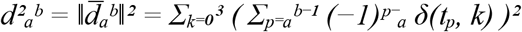

##### Distance penalty 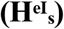

Compels contact residues to remain at a physical lattice distance of

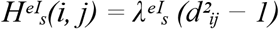

where 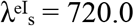 is the distance penalty scale factor.

##### Locality penalty (H_lo_C)

Enforces self-avoiding local packing around the active contact point to prevent steric overlap:

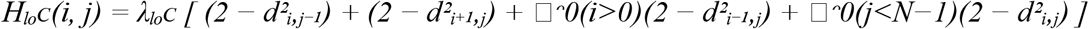

where λ_lo_C = 20.0 is the locality penalty coefficient and □ᵔ0 is the indicator function.

### 2.3. Gate Based Variational Quantum Eigensolver with CVaR

In the gate-based VQE framework, the total quantum register is initialised with 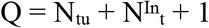 qubits, where N_tu_ represents the active configuration qubits, 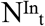 the interaction control qubits, and the remaining qubit serves as an entangling ancilla.

#### 2.3.1 Parameterised Ansatz Design

The quantum state |ψ(θ)⟩ is prepared by applying a parameterised unitary ansatz U(θ) to the vacuum state:

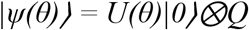

The scalable hardware-efficient ansatz U(θ) is structured as follows:

1. **Superposition layer:** An initial layer of Hadamard gates on all qubits, H⊗Q.
2. **First rotation layer:** Parameterised R_γ_(θ_k_) rotation gates applied to target qubits 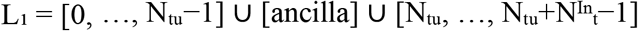.
3. **Entangling CX loop:** A circular CNOT chain building quantum correlations across all degrees of freedom: a forward chain across configuration qubits, CNOT from the final configuration qubit to the ancilla, CNOT from the ancilla to the final interaction qubit, a reverse chain across interaction qubits, and a closing CNOT from the first interaction qubit back to the first configuration qubit.
4. **Second rotation layer:** Parameterised R_γ_(θ_1_) rotation gates applied to target qubits 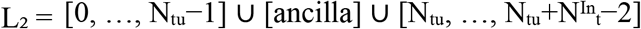.

Measurements are performed exclusively on the 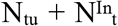 computational qubits, tracing out the ancilla.

#### 2.3.2 Conditional Value at Risk (CVaR) Objective

To mitigate NISQ-era gate errors, decoherence, and sampling variance, the optimisation loop employs a Conditional Value at Risk (CVaR) objective. For a tail parameter α ∈ (0, 1], CVaRα(H) is defined as the expected energy over the lowest α percentile of the measured state distribution:

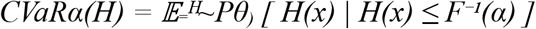

where Pθ(x) = |⟨x|ψ(θ)⟩|^2^ and F(e) = Pθ(H(x) ≤ e) is the CDF of the energy. For M discrete bitstrings sorted by energy E_1_ ≤ E_2_ ≤ ⋯ ≤ E^M^ with probabilities p_1_, p_2_, …, p^M^, the CVaR objective evaluates as:

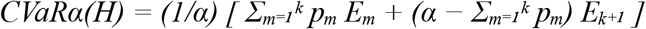

where k is the largest index satisfying 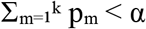.

#### 2.3.3 Classical Optimisation

The parameters θ are optimised classically using the Nelder–Mead simplex algorithm. To circumvent local minima in the rugged energy landscape, a multi-start heuristic executes 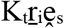 independent optimisation runs with parameters randomly initialised from a uniform distribution θ_j_ ~ *U*(−π, π). The run achieving the minimum CVaR energy is selected as the final solution.

### 2.4. Quantum Annealing Formulation and QUBO Translation

To solve the peptide folding problem on superconducting quantum annealers (e.g., D-Wave Advantage), the global Hamiltonian is mapped to a Quadratic Unconstrained Binary Optimisation (QUBO) form:

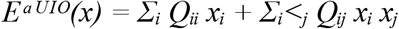

where x_i_ ∈ {0, 1} represent the combined set of configuration, interaction, and auxiliary binary variables.

#### 2.4.1 Degree Reduction of High Order Terms

Because the distance metrics 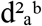 involve products of Kronecker delta terms, they introduce quartic (degree 4) terms in the configuration variables. The interaction Hamiltonian further multiplies contact control variables c_ij_ by these distance metrics, yielding quintic (degree 5) polynomial terms of the form 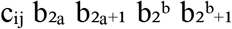. High-order terms are reduced to quadratic form by recursively introducing auxiliary binary variables. For any product z = x_1_x_2_, we substitute a new binary variable z and enforce the equivalence via the penalty:

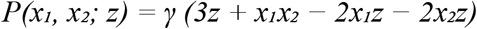

where γ > 0 is a penalty scaling factor chosen such that γ > max(0, −E^epn^). This penalty is minimised (P = 0) if and only if z = x_1_x_2_. Chaining this reduction, a degree d term is translated into quadratic terms with the addition of d−2 auxiliary variables. The QUBO degree-reduction schematic is provided as supplementary figure 1.

#### 2.4.2 Minor Embedding and Annealing Schedule

The resulting sparse quadratic coupling matrix Q is mapped onto the physical hardware graph (e.g., Pegasus topology) using heuristic minor embedding algorithms. To preserve the coherence of logical variables mapped to physical qubit chains, the chain coupling strength 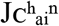 is tuned relative to the maximum elements of Q:

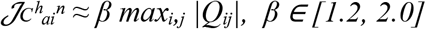

The system is initialised in the uniform superposition state and evolved according to the time-dependent transverse-field Ising Hamiltonian:

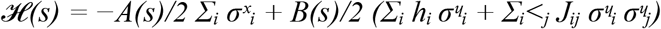

where s = t/τ ∈ [0, 1] is the normalised time, τ = 20 μs is the total annealing duration, and A(s), B(s) are the annealing envelope functions. The linear biases h_i_ and quadratic couplings J_ij_ are obtained from the QUBO parameters Q via the standard transformation 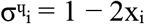.

### 2.5. Simulated Annealing Protocol

Simulated Annealing (SA) is employed as a classical metaheuristic to search the discrete peptide folding energy landscape. The algorithm uses a Markov Chain Monte Carlo (MCMC) transition kernel to escape local minima. The full search space is represented as a joint binary state vector:

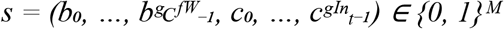

where 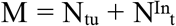 is the total dimensionality. The target is to locate the global minimum s* = argmin_s_ H(s).

#### 2.5.1 Initialisation

An initial configuration s^−0(^ ∈ {0, 1}^M^ is selected either randomly or from a coarse-grained secondary structure heuristic. The initial temperature T_0_ is set such that almost all transitions are initially accepted, with a cooling parameter γ ∈ (0.95, 0.999), minimum temperature T^epn^ *≪* 1, and per temperature step budget L.

#### 2.5.2 Neighbourhood Proposal and Metropolis–Hastings Acceptance

At each step, a candidate state s’ is generated by selecting an index j ∈ {0, …, M−1} uniformly at random and flipping the bit: s’_j_ = 1 − s_j_; s’^e^ = s^e^ for all m ≠ j. The energy difference ΔE = H(s’) − H(s) is computed exactly, and the candidate is accepted with the Boltzmann probability:

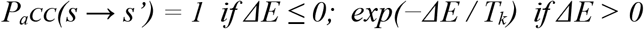

#### 2.5.3 Cooling Schedule and Termination

After L transition attempts at temperature T_k_, the temperature is decreased geometrically: T_k+1_ = γT_k_. The algorithm terminates when T_k_ < T^epn^. The best state s* is then mapped to 3D backbone Cartesian coordinates using the discrete dihedral angle mapping (TURN_DIHEDRAL).

### 2.6. Hybrid SA–VQE Decomposition Pipeline

To scale the simulation to longer peptides without exceeding physical qubit limits on NISQ hardware, a hybrid decomposition strategy combines SA and VQE across two sequential phases:

The total search space is split into the discrete interaction contact topology 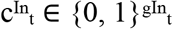 and the active backbone configuration angles 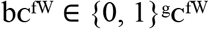.

SA is run on the full Hamiltonian to identify the most stable contact topologies. The contact variables are frozen to their optimal values: 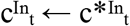.

The VQE Hamiltonian is projected onto the subspace where interaction variables are fixed, eliminating 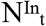 qubits from the quantum register and reducing the required qubit count to 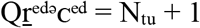.

Gate-based VQE searches the reduced state space to optimise the backbone configuration angles bC^fW^ under the static contact topology 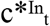.

The combined bitstring (b*C^fW^, 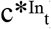) is mapped to 3D physical coordinates via bond lengths and dihedral angles (NeRF-like reconstruction) and exported as a PDB structure. This is provided as a mermaid diagram in (Fig 1B).

### 2.7. Discrete 2D Square Lattice Representation

During both classical search (Simulated Annealing) and quantum optimisation (VQE), peptide conformation is modelled as a self-avoiding walk on a 2D square lattice. Each turn code t_a_ encodes a relative heading change with respect to the current direction of travel, providing a computationally compact representation of backbone geometry prior to 3D reconstruction.

#### 2.7.1 Heading Update and Coordinate Tracing

The walk begins at position 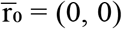 with an initial heading vector 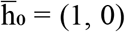 (facing +x). The four turn codes correspond to the following heading changes:

- **t**_**a**_ **= 0 → Straig ht** (no heading change; heading is +x relative to current direction)
- **t**_**a**_ **= 1 → Left Turn** (+90° counterclockwise)
- **t**_**a**_ **= 2 → Right Turn** (−90° clockwise)
- **t**_**a**_ **= 3 → Reverse** (180° turn; heavily penalised)

For each residue a ∈ {0, …, N−2}, the heading 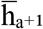 is updated dynamically based on the turn t_a_:

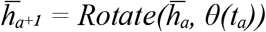

where the turn angles are θ(0) = 0, θ(1) = π/2, θ(2) = −π/2, and θ(3) = π. The 2D lattice coordinates of residue a+1 are then:

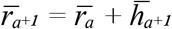

#### 2.7.2 Dihedral Angle Mapping for 3D Reconstruction

For PDB export, the discrete turn codes are mapped to continuous backbone dihedral angles (φ_a_, ψ_a_) approximating standard secondary structural motifs. Table 1 summarises the complete mapping.

**Table 1.** Turn code to backbone dihedral angle mapping.

| Turn Code<br>($t_a$) | Binary<br>($b_{2a}b_{2a+1}$) | Dihedral Angles ($\phi_a, \psi_a$) | Structural Motif |
| --- | --- | --- | --- |
| 0 | 00 | (-60°, -45°) | Helix-like conformation |
| 1 | 01 | (-135°, 135°) | $\beta$ strand-like conformation |
| 2 | 10 | (-75°, 145°) | Polyproline II-like conformation |
| 3 | 11 | (-60°, 140°) | Extended/coil-like conformation |

### 2.8. Overlap Penalty Term

The updated total conformational energy Hamiltonian includes a dedicated overlap penalty term that prevents lattice self-intersection:

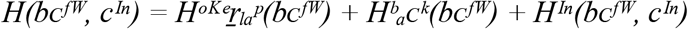

Lattice self-intersection is prevented by applying a severe overlap penalty. If any two distinct residues i and j occupy the same coordinate on the 2D lattice (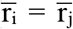 for i ≠ j), the energy is penalised:

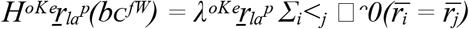

where 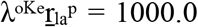 by default. In the precomputed energy lookup tables, any configuration yielding 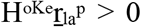 is classified as a forbidden state and assigned energy ∞ to stabilise both classical and quantum optimisations.

### 2.9. Parameter Tuning Rationale

Backbone penalty 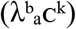 : In early implementations, 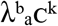 was set to 50.0 to enforce strong backbone diversity. In the current package configuration, this value has been reduced to **5.0** so that Miyazawa–Jernigan contact energies (typically ranging from −4.0 to +4.0) dominate the objective landscape, allowing more nuanced contact-driven folding.

Distance and locality penalties 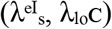 : Both penalties are **disabled by default** 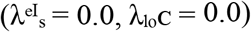 in the modern package. In earlier formulations, these were set to 720.0 and 20.0 respectively; although gated by the contact control bits c_ij_, large penalty values could suppress the discovery of favorable folding contacts by incentivizing the optimizer to set c_ij_ = 0. These weights can be re enabled for specific experimental runs and are applied conditionally on lattice contacts.

### 2.10. Modular Hybrid Pipeline

The modern QuPepFold package expands the hybrid SA–VQE decomposition (described in §6) into a fully modular 8 stage pipeline designed to scale to longer peptides while remaining compatible with NISQ hardware constraints.

#### Stage 1 Global Simulated Annealing

A classical SA search is executed on the full chain with 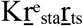 random restarts (default: 3) to locate a high-quality global scaffold. This stage yields the best global turns vector 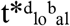, positions 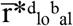, and energy 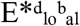. Temperature follows a geometric schedule: T_k+1_ = γT_k_ with γ ∈ [0.95, 0.999].

#### Stage 2 Fragment Generation

The full sequence is divided into overlapping fragments of length 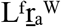 (default: 7 amino acids) with an overlap of 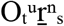 turns (default: 2). For each fragment i, the start residue index S_i_ and end residue index E_i_ are computed to ensure full sequence coverage.

#### Stage 3 Energy Table Precomputation

For each fragment (e.g., length 7 has 5 turns, requiring 2×5 = 10 bits), all 2^10^ = 1024 local conformational states are enumerated. Self-intersecting walks are assigned ∞ energy. The Miyazawa–Jernigan potential is evaluated for all contacting residue pairs within the fragment, forming a lookup table used by the subsequent VQE stage.

#### Stage 4 Quantum VQE Refinement with QA Anchored Optimisation

Each fragment is refined independently using a local VQE loop. To preserve global structural coherence, QuPepFold employs QA Anchored Optimisation:

- **Boundary freezing:** Turn bits overlapping with adjacent fragments are fixed to the values from the global SA solution 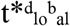. Any state violating this boundary constraint is assigned ∞ energy in the local optimisation table.
- **Context energy evaluation:** The fragment is aligned to the global coordinate framework. The interaction energy between the fragment’s active residues and the fixed global environment is added to the local Hamiltonian:

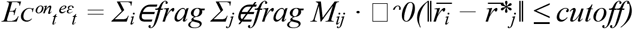

where 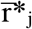 are the fixed coordinate positions of the global environment.

#### Stage 5 Fragment Stitching via Dynamic Programming

VQE-derived fragment candidates are assembled into a global conformation using a Shortest Path Dynamic Programming (DP) algorithm. Let dp[i][k] denote the minimum energy to reach candidate k of fragment i:

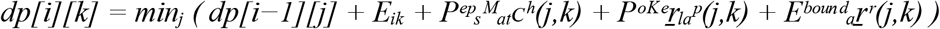

where 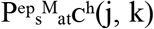(j, k) penalises mismatched bits in the overlap region (set to ∞ in strict mode or a soft weight in relaxed mode); 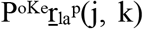 (j, k) penalises combined paths that produce lattice self-intersections; and 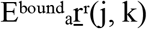 (j, k) captures cross-fragment contact energy between boundary residues of fragments i−1 and i.

#### Stage 6 3D Coordinate Rebuilding

The best stitched bitstring is decoded into turn codes and backbone Cartesian coordinates are reconstructed recursively using a NeRF-like (Natural Extension Reference Frame) placement algorithm, starting from a seeded first residue 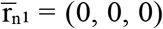 and applying standard peptide bond lengths, angles, and turn dihedral mappings. Cβ atoms are placed using tetrahedral geometry for all non-glycine residues.

#### Stage 7 Output Reporting

Results are compiled into the output directory as described in §7.3, supplemented by the fragment energy tables and DP stitching trace where applicable.

### 2.11. Local VQE Ansatz and Biased Warm Start Initialisation

#### 2.11.1 Modular Fragment Level Ansatz

For fragment refinement in Stage 4, QuPepFold employs a shallower, hardware-efficient ansatz of depth D over 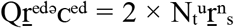 qubits (two configuration qubits per turn). The ansatz structure is:

**Hadamard layer:**Superposition on all active qubits.

**R**_**γ**_ **rotation layer:**Parameterized R_γ_(θ^d^,_i_) gates on each qubit i at depth d.

**Linear CNOT chain:**CNOT gates applied sequentially: 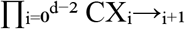.

**Loop closure:**On the final layer (d = D−1), an additional loop closing CNOT is applied between the last and first qubit: 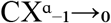.

#### 2.11.2 Biased R_γ_ Warm Start Initialization

To accelerate convergence, QuPepFold can warm start the local VQE state preparation using the classical Simulated Annealing solution. If a target bitstring z_0_ is provided, each qubit is initialised into a biased superposition prior to the ansatz using custom R_γ_ rotations:

- **If target bit z**_**0**_,_**i**_ **= 1:** apply R_γ_(θ_1_, i) where θ_1_ = 2 arcsin(√β).
- **If target bit z**_**0**_,_**i**_ **= 0:** apply R_γ_(θ_0_, i) where θ_0_ = 2 arcsin(√(1−β)).

Here, β ∈ (0.5, 1) is the warm start bias (default: 0.95). When measured, each qubit yields the correct classical bit value with probability β, seeding the variational optimisation close to the classical solution.

### 2.12. CVaR Objective with Forbidden State Filtering

The CVaR objective function is augmented with an explicit forbidden state filtering step to handle self-intersecting configurations. The full computation proceeds as follows:

All configurations with ∞ energy (self-intersecting walks or boundary violations) are excluded from the state distribution.

The remaining probability values are renormalised over the set of valid states:

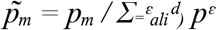

States are sorted by energy E_1_ ≤ E_2_ ≤ ⋯ ≤ E^P^. The CVaR objective is then evaluated over the α tail of the renormalised distribution:

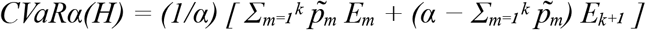

where k is the largest index satisfying 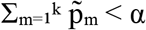.

### 2.13. Ising Coefficient Conversion from QUBO

When translating the QUBO problem to the Ising formulation required by physical quantum annealers, the linear biases h_i_ and quadratic couplings J_ij_ are derived from the QUBO coefficient matrix Q via the standard transformation 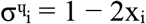 :

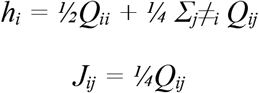

These coefficients are passed directly to the annealer’s programming interface. The chain coupling strength for minor embedded logical qubits is set as 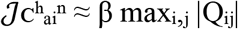, with β ∈ [1.2, 2.0] tuned empirically to balance chain integrity against problem energy scale.

### 2.14. Configuration Parameters

Table 2 provides a complete listing of all configuration parameters in the FoldConfig specification, including parameters added in the QuPepFold > v1.0.0.

**Table 2.**
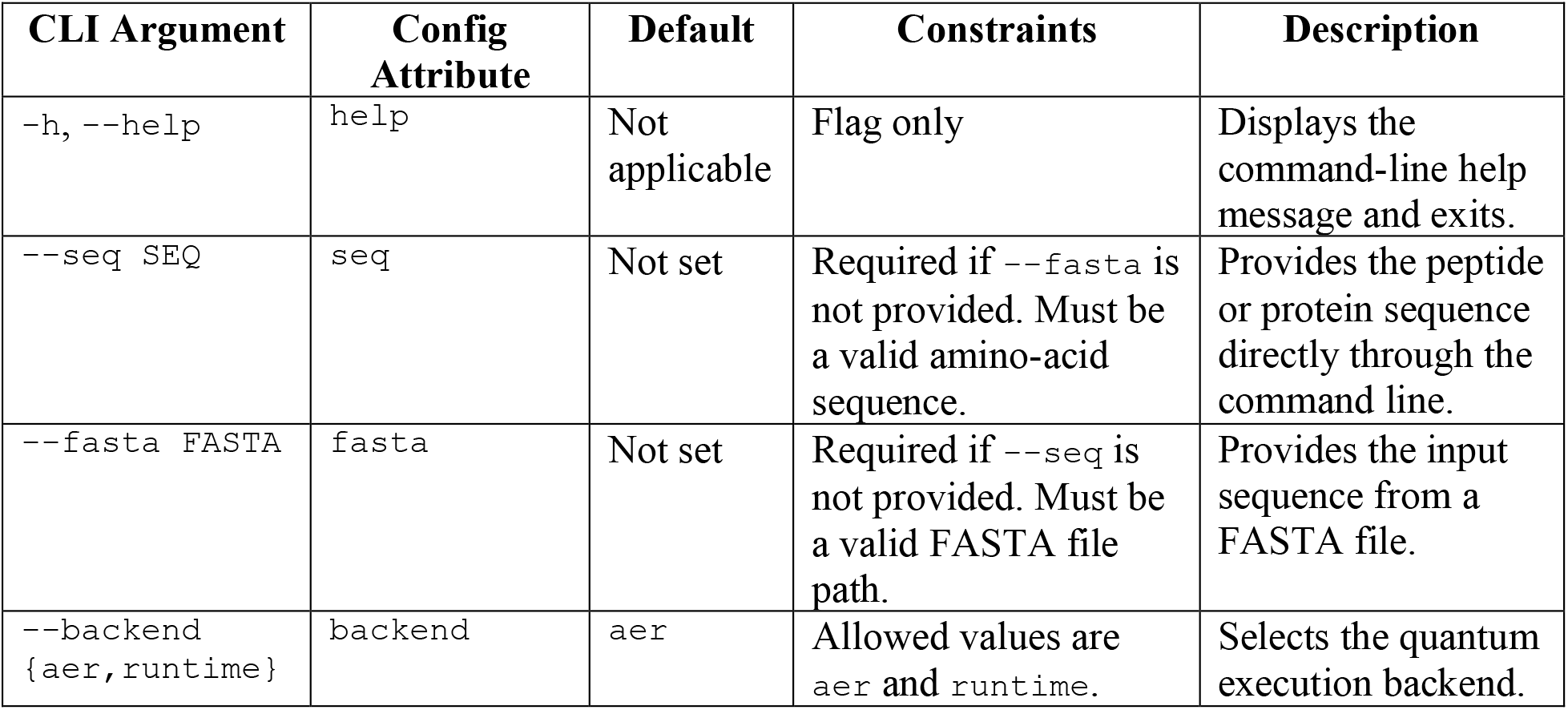
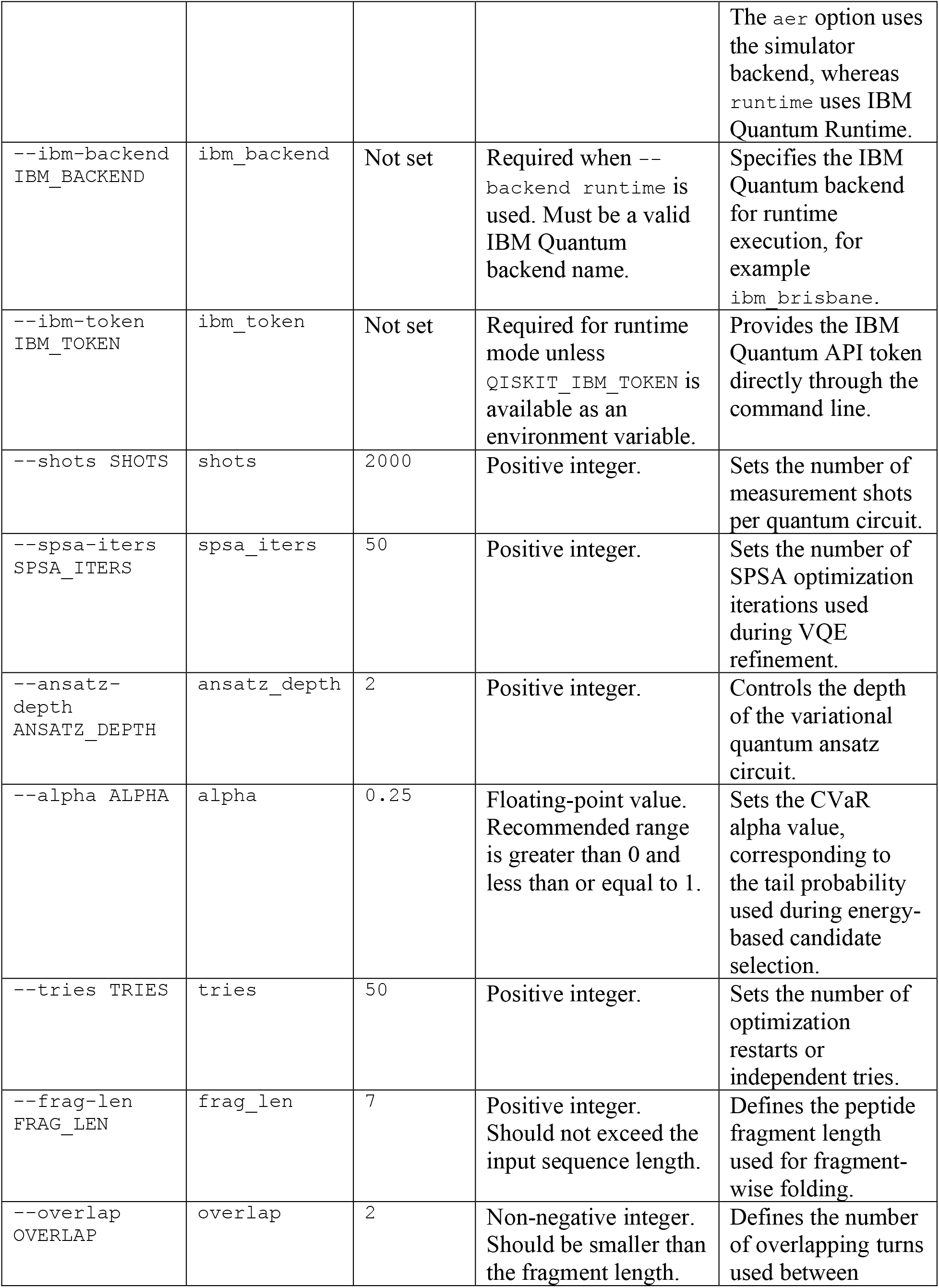
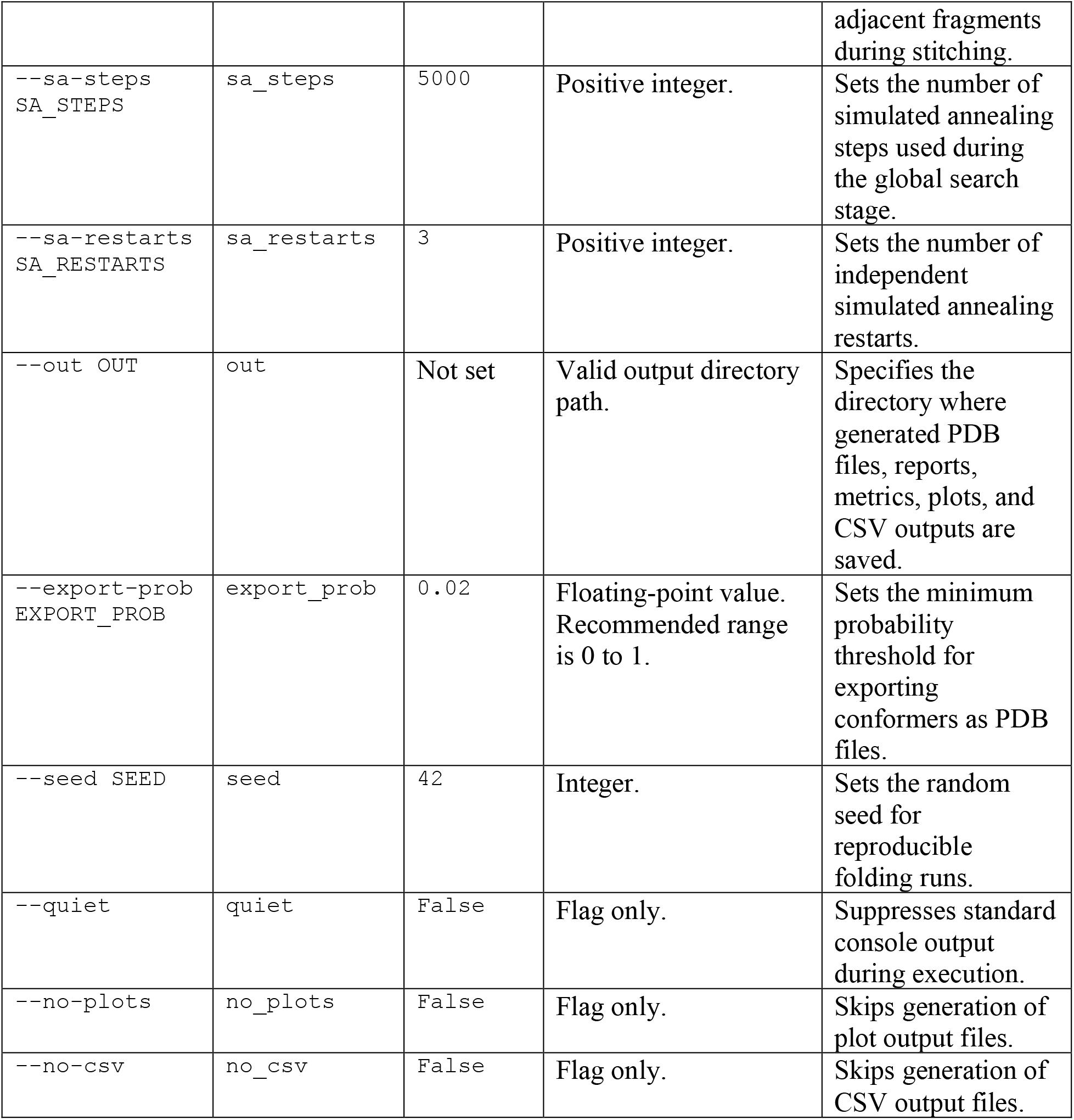
Complete QuPepFold configuration parameters.

## 3. Results

The quantum-classical folding workflow implemented was evaluated on the 13-residue peptide ETFSDLWKLLPEN, corresponding to the reference segment extracted from the 1YCR crystal structure. Using the exact matching sequence removes any ambiguity from sequence-length differences, allowing direct structural comparison between the predicted and reference conformations. Folding was performed using ibm_fez with 2,048 shots. Hybrid VQE refinement and relaxed stitching were enabled. The pipeline was configured for 5,000 simulated annealing (SA) steps across three independent restarts, 50 SPSA iterations, a fragment length of seven residues, and two overlapping turns between adjacent fragments. Three independent simulated annealing runs produced internal energies of −3.46, −3.53, and −3.70, respectively, as shown in (Fig 2A). The third run yielded the most favorable conformation, and the corresponding bitstring was selected as the leading global solution. All three runs converged to negative-energy conformers, confirming that the global search covered the landscape adequately for this 13-residue sequence.

**Figure 2:**
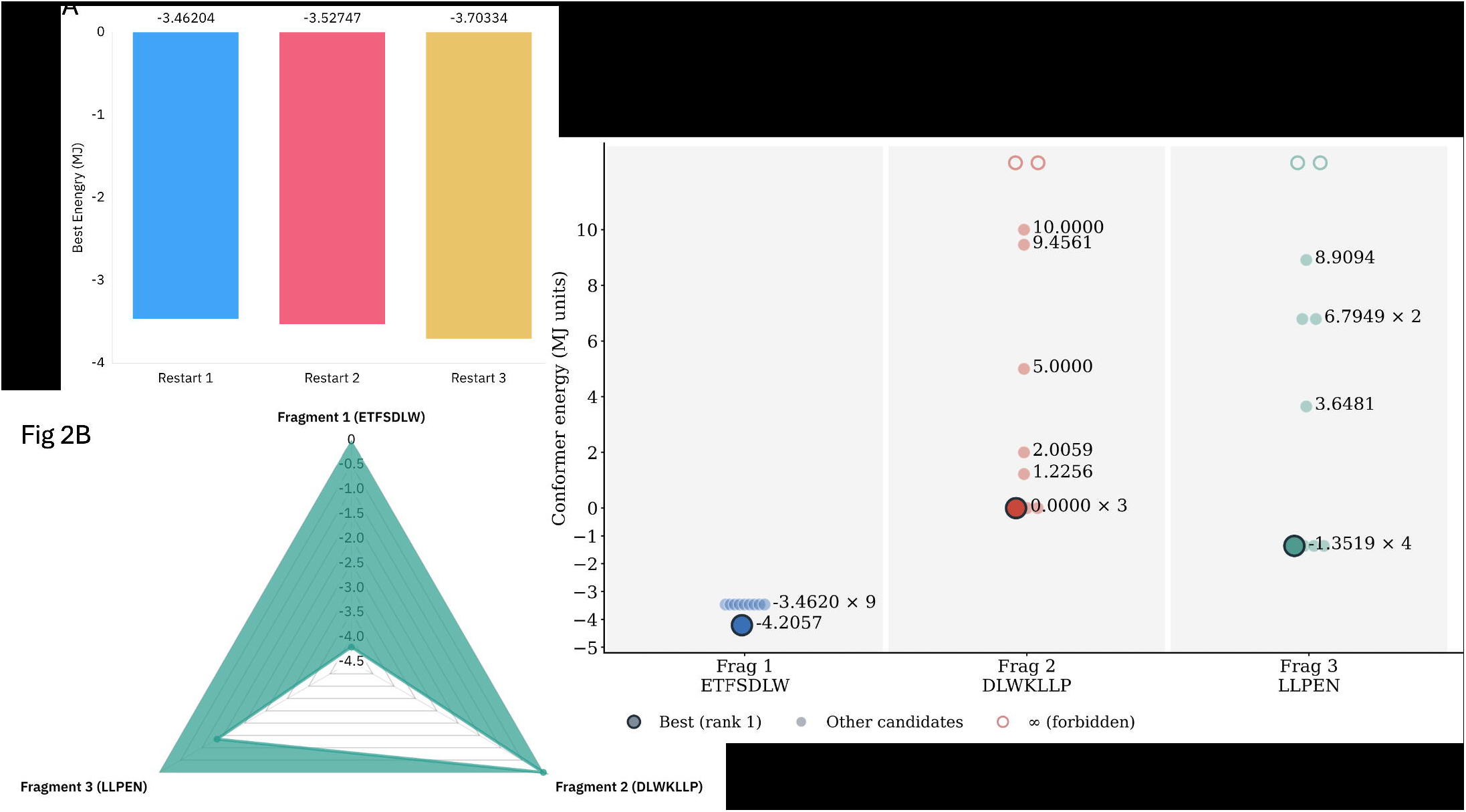
Energy landscape comparison. A) Energy trace (y: conformational energy, x: SA step) for all three restarts. B) Best conformer energy returned by VQE per fragment. Frag 1 (ETFSDLW): −4.21 MJ. Frag 2 (DLWKLLP): E = 0.00 (no favourable contacts). Frag 3 (LLPEN): −1.35 MJ. C) Three panels (one per fragment: ETFSDLW, DLWKLLP, LLPEN) with scatter plot of energy across the 10 VQE candidate conformers, with the selected candidate (minimum energy) highlighted.

The peptide was partitioned into three overlapping fragments for local quantum-classical refinement: `ETFSDLW` (residues 0–7), `DLWKLLP` (residues 4–11), and `LLPEN` (residues 8–13). An energy table 3 of 1,024 possible states was constructed for each fragment. VQE refinement generated 10 candidate conformational states per fragment, producing 30 candidates in total.

**Table 3.** Fragment-level VQE energetics for the three overlapping peptide fragments.

| Fragment | Residues | SPSA Energy | Best Candidate | Status |
| --- | --- | --- | --- | --- |
| ETFSDLW (N-terminal) | 0–7 | $-4.21$ | $-4.21$ | Favorable |
| DLWKLLP (Central) | 4–11 | 1000.00 | 0.00 | Poor |
| LLPEN (C-terminal) | 8–13 | $-0.73$ | $-1.35$ | Favorable |

Across all VQE-generated candidates, 14 fragment conformers with negative energies were identified. The N-terminal ETFSDLW fragment produced the most favorable local result (SPSA energy = −4.21; best candidate = −4.21), contributing the majority of negative-energy states. The C-terminal LLPEN fragment also yielded negative-energy candidates (SPSA energy = −0.73; best candidate = −1.35). In contrast, the central DLWKLLP fragment showed anomalous optimization behavior, with a reported SPSA energy of 1,000.00 and a best candidate energy of 0.00, indicating that the local VQE search failed to identify favorable conformers for this segment.

The poor performance of the central fragment may reflect proline’s restricted backbone torsion angles, though the anomalous energy value (1,000.00) could equally indicate a failure in scoring, encoding, or the search procedure rather than an intrinsic chemical limitation.

Fragment stitching was performed using dynamic programming under the *strict* stitching mode. The detailed results are tabulated in supplementary table 1. The selected states carried energies of −4.21 (ETFSDLW), 0.00 (DLWKLLP), and −1.35 (LLPEN) as shown in (Fig 2B). The initial stitched chain exhibited an energy of +8.73, reflecting incompatibilities introduced when independently optimized fragments were assembled into a contiguous chain. Fragment-level individual values of 10 VQE candidate conformers are shown in (Fig 2C).

The MJ matrix interaction energy is shown in Fig 3.

**Figure 3:**
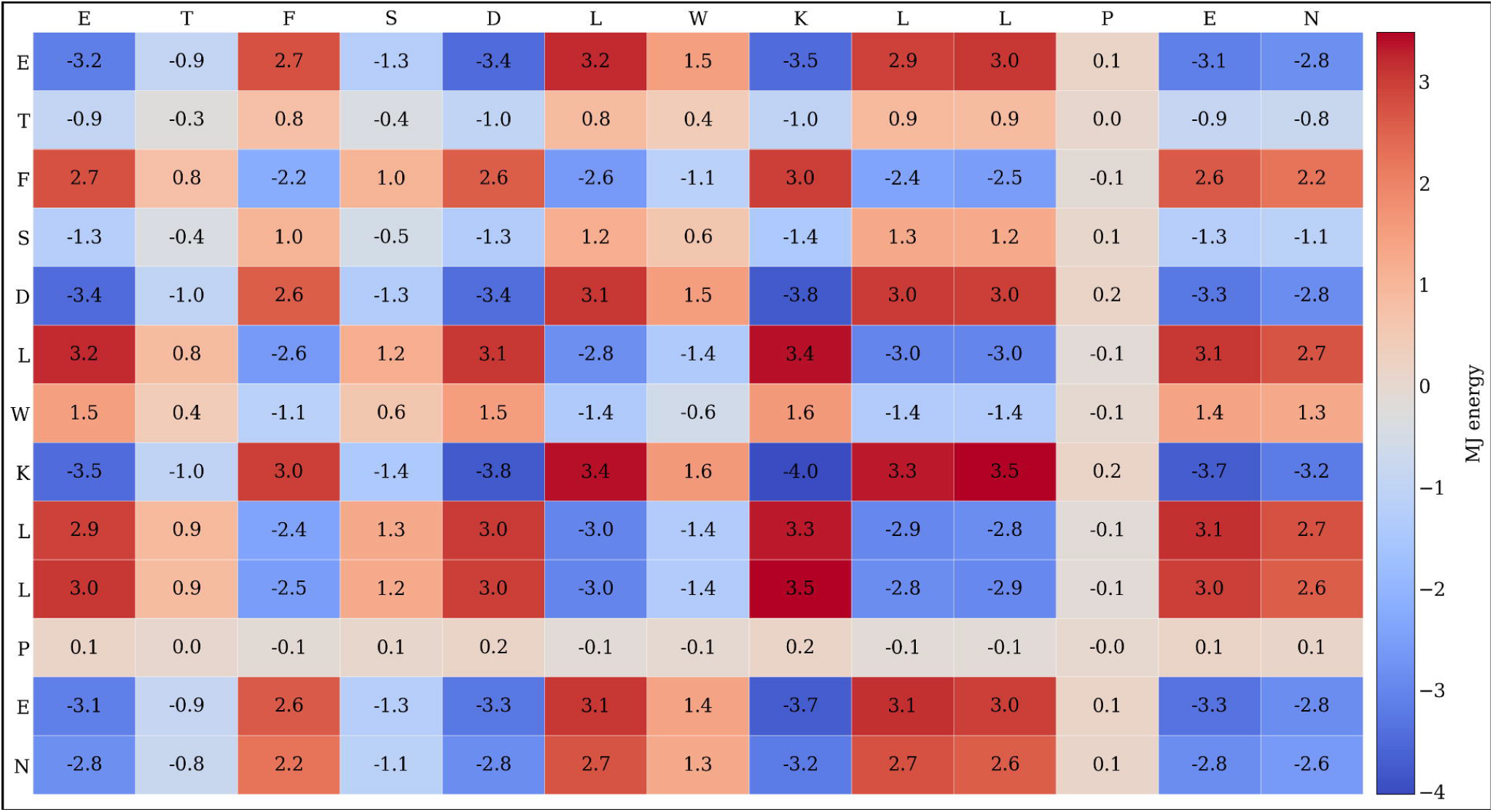
Miyazawa–Jernigan (MJ) pairwise contact energy matrix. Residue–residue MJ statistical potential values (MJ units). Negative (blue) = favourable; positive (red) = unfavorable.

Post-stitch refinement brought the chain energy from +8.73 to −2.58, but this still fell short of the global SA result (−3.70). QuPepFold therefore selected the global SA solution as the final output. The final backbone-building stage generated a 65-atom model and wrote the predicted structure to a PDB file.

This outcome means the 3D backbone model reported in Fig. 4 and the RMSD values in Table 4 are derived exclusively from the classical simulated annealing stage. The VQE refinement and dynamic-programming stitching pipeline produced a valid conformer (E = −2.58 MJ) but did not surpass the global SA solution in this run. We emphasise that this is expected behaviour in the current NISQ regime: VQE with 50 SPSA iterations on a noisy simulator cannot guarantee improvement over a multi-restart SA search with 5,000 steps. The contribution of the quantum component here is therefore not a better final structure, but (i) generation of a ranked library of 30 fragment-level candidates with associated sampling probabilities, (ii) a fragment-stitched conformer at −2.58 MJ that is independently valid, and (iii) a complete end-to-end workflow demonstration that is auditable and reproducible.

**Table 4.** Structural comparison of the QuPepFold model versus the 1YCR reference peptide.

| Analysis | Atom Pairs | RMSD (Å) | Interpretation |
| --- | --- | --- | --- |
| Full backbone superposition | 38 | 4.6456 | Poor global agreement |
| Trimmed alignment (10 residues) | — | 2.90 | Partial local similarity |

**Figure 4:**
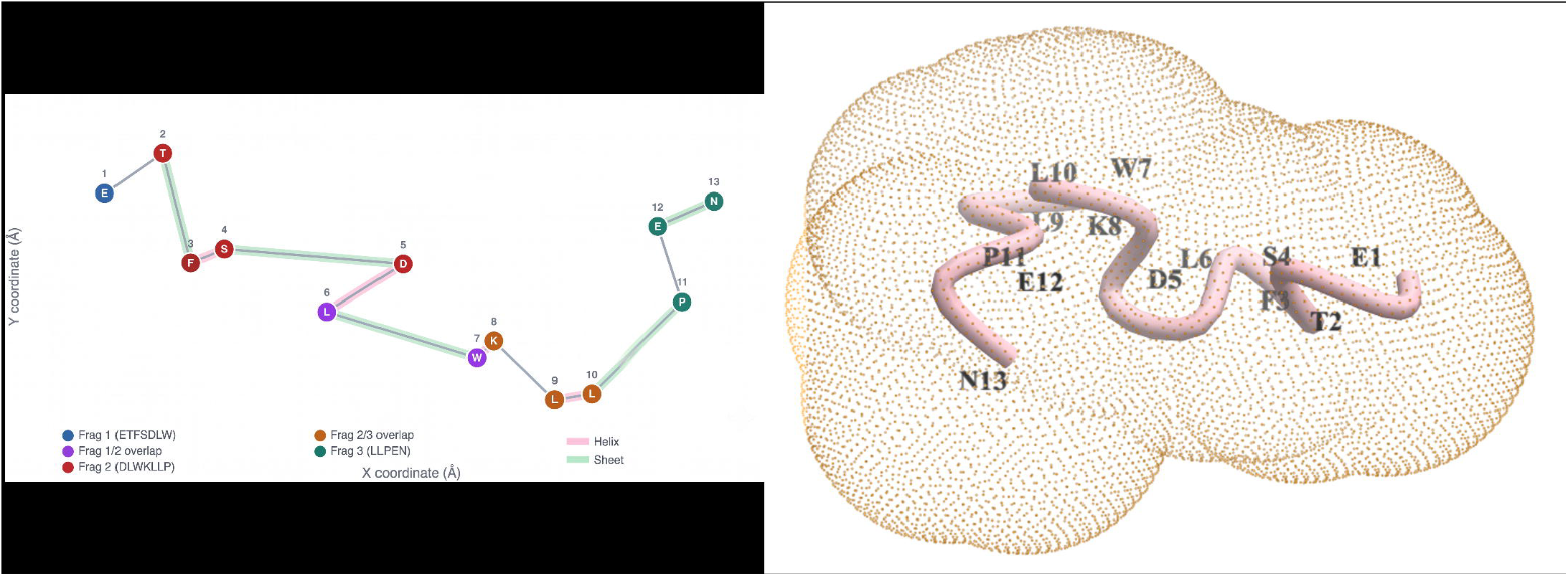
Predicted backbone C_α_ trace. A) 2D projection (X–Y) of 13-residue backbone Cα positions from backbone.pdb. Residues coloured by fragment assignment. Shaded spans: pink = helix, green = sheet. B) The 3D visualization of backbone PDB structures generated. These are visualized as ribbon with dot mesh overlaid.

The backbone showed helix-like geometry at residue pairs 3–4, 5–6, 7–8, and 9–10, and sheet-like geometry at pairs 2–3, 4–5, 6–7, 10–11, and 12–13. These two-residue annotations do not constitute canonical secondary structure elements; they describe local backbone geometry only. This is depicted as a 2D overlay in (Fig 4A). The 3D fold of the sequence is shown in (Fig 4B).

Structural comparison against the PDB reference used ICM with backbone provided in Table 4 for N, Cα, and C atoms — a stricter criterion than Cα-only alignment. One atom pair was excluded due to zero occupancy, leaving 38 pairs for RMSD calculation.

The full-backbone RMSD of 4.65 Å indicates poor global structural agreement. Because the sequences are identical, this cannot be explained by mismatch; it reflects a genuine difference in backbone geometry. For a 13-residue peptide, 4.65 Å is substantial and does not support a claim of global fold recovery. The values are depicted in (Fig 5A and Fig 5B). A trimmed 10-residue alignment improved the RMSD to 2.90 Å, showing that part of the backbone has moderate local accuracy. The gap between full (4.65 Å) and trimmed (2.90 Å) values points to localised regions of poor fit alongside segments of reasonable agreement.

**Figure 5:**
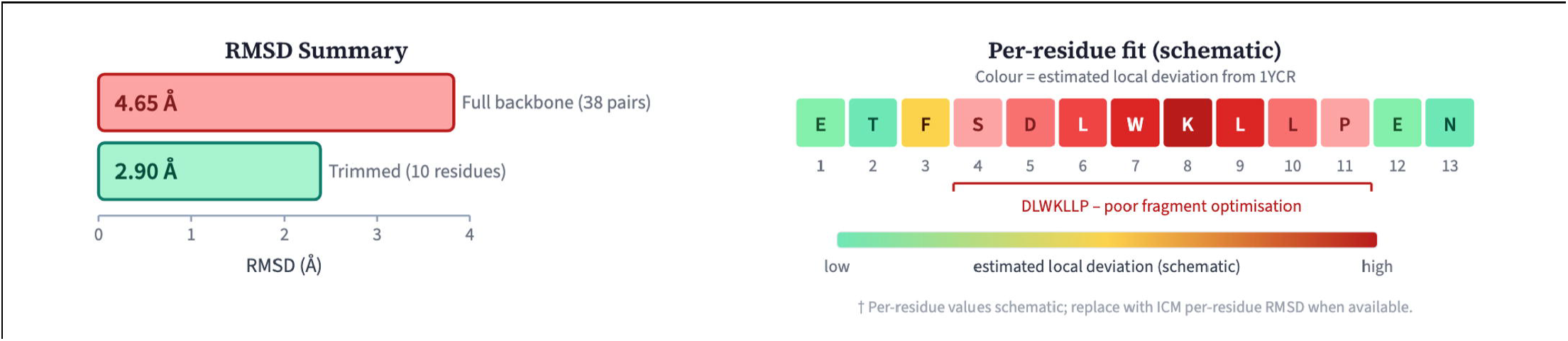
Backbone RMSD comparison. A) Full backbone (38 pairs, RMSD = 4.65 Å) vs. trimmed 10-residue alignment (RMSD = 2.90 Å). B) Per-residue schematic coloured by estimated local deviation.

## 4. Discussion

QuPepFold enters a field in which quantum and hybrid quantum–classical solvers are being applied to the protein folding problem with increasing methodological sophistication. The energetic and structural results reported here—successful generation of a complete 65-atom backbone model for the p53 transactivation peptide ETFSDLWKLLPEN, identification of a negative-energy global conformer (E = −3.70), but a full-backbone RMSD of 4.65 Å against the 1YCR reference—are most usefully interpreted not in isolation but against the broader landscape of classical, quantum-annealing, and gate-based variational approaches to structure prediction. The sections below compare QuPepFold’s methodological choices against the relevant literature, identify where the results align with or depart from prior work, and define the conditions under which quantum-enabled folding is currently meaningful.

### 4.1 Relationship to the classical structure-prediction paradigm

Any quantum approach to folding must ultimately be measured against the classical state of the art. The thermodynamic hypothesis underpinning all of these methods traces to Anfinsen’s demonstration that the native conformation is encoded in the primary sequence and corresponds to a free-energy minimum [1], while the combinatorial vastness that motivates non-deterministic search was famously articulated by Levinthal [2]. Classical physics-based sampling—molecular dynamics and Monte Carlo with empirical force fields—remains limited by the femtosecond integration step relative to the millisecond-to-second timescales of biological folding, and by entrapment in local minima of a rugged landscape [3,4]. Knowledge-based fragment-assembly methods such as Rosetta circumvented brute-force sampling by recombining experimentally observed fragments under statistical potentials, and remain a strong baseline for short peptides and loops. The more consequential advance came from deep learning: AlphaFold2 demonstrated near-experimental accuracy for the majority of single-domain proteins at CASP14 by learning evolutionary and geometric constraints directly from the structure database, and RoseTTAFold [14,15] reproduced much of this performance with a three-track architecture.

It is essential to position QuPepFold honestly against this backdrop. For a well-behaved globular domain with deep multiple-sequence alignments, a co-evolution-driven predictor will currently outperform any near-term quantum method by a wide margin, and the present 4.65 Å backbone RMSD on a 13-residue peptide underscores that quantum-enabled folding is not yet competitive on a head-to-head accuracy basis. The legitimate niche, rather, lies precisely where the data-driven paradigm is weakest: short, conformationally plastic peptides and intrinsically disordered regions (IDRs) that lack a single dominant fold, shallow alignments, and the ensemble character of binding-competent states. QuPepFold’s explicit targeting of IDRs for therapeutic exploration is therefore a deliberate and defensible scoping choice rather than a claim to supersede AlphaFold-class predictors [16].

### 4.2 Quantum annealing versus gate-based variational formulations

Quantum annealing and gate-based variational eigensolvers represent the two main hardware paradigms for folding, each with distinct trade-offs that QuPepFold addresses through its dual QUBO and VQE formulations. The earliest quantum-hardware demonstration of lattice folding was the quantum-annealing study of Perdomo-Ortiz and co-workers, who mapped an HP lattice model to a QUBO and solved it on early D-Wave hardware [17,18]. That lineage continues in subsequent HP-lattice annealing work and in D-Wave’s own lattice-protein design demonstrations on the Pegasus topology [19-22]. Annealing offers large physical qubit counts and a natural fit for QUBO-encoded Ising problems, but its solution quality is constrained by limited connectivity—necessitating minor embedding and chain-coupling tuning, exactly the J_chain calibration described in our QUBO translation—and by the absence of programmable coherent gate operations. The degree-reduction strategy QuPepFold uses to collapse the quintic terms arising from the distance metrics into quadratic form with auxiliary variables is the same class of penalty-based gadget reduction that all QUBO folding formulations must employ, and it inflates qubit count substantially, which is the central practical limitation of the annealing route.

Gate-based VQE, by contrast, exploits programmable entanglement and a tunable ansatz, and was placed on a rigorous resource-efficient footing by Robert and colleagues, who folded the 10-residue Angiotensin peptide on a tetrahedral lattice while compressing the register from 35 to 22 qubits [24]. QuPepFold inherits this resource-efficient philosophy and extends it: the hybrid SA–VQE decomposition freezes the contact topology c* obtained from a classical pass and projects the VQE Hamiltonian onto the reduced subspace Q_reduced = N_cfg + 1, removing the entire interaction register from the quantum circuit. This is conceptually aligned with the broader consensus, articulated in the perspective of Babbush and co-workers, that near-term advantage is far more likely from hybrid decomposition than from monolithic quantum solution of the full problem [4,5], and with the wider survey of variational quantum algorithms for chemistry and drug discovery [10]. This reduction in qubits is depicted in (Fig 6A). The present work therefore follows the mainstream methodological direction of the field.

**Figure 6:**
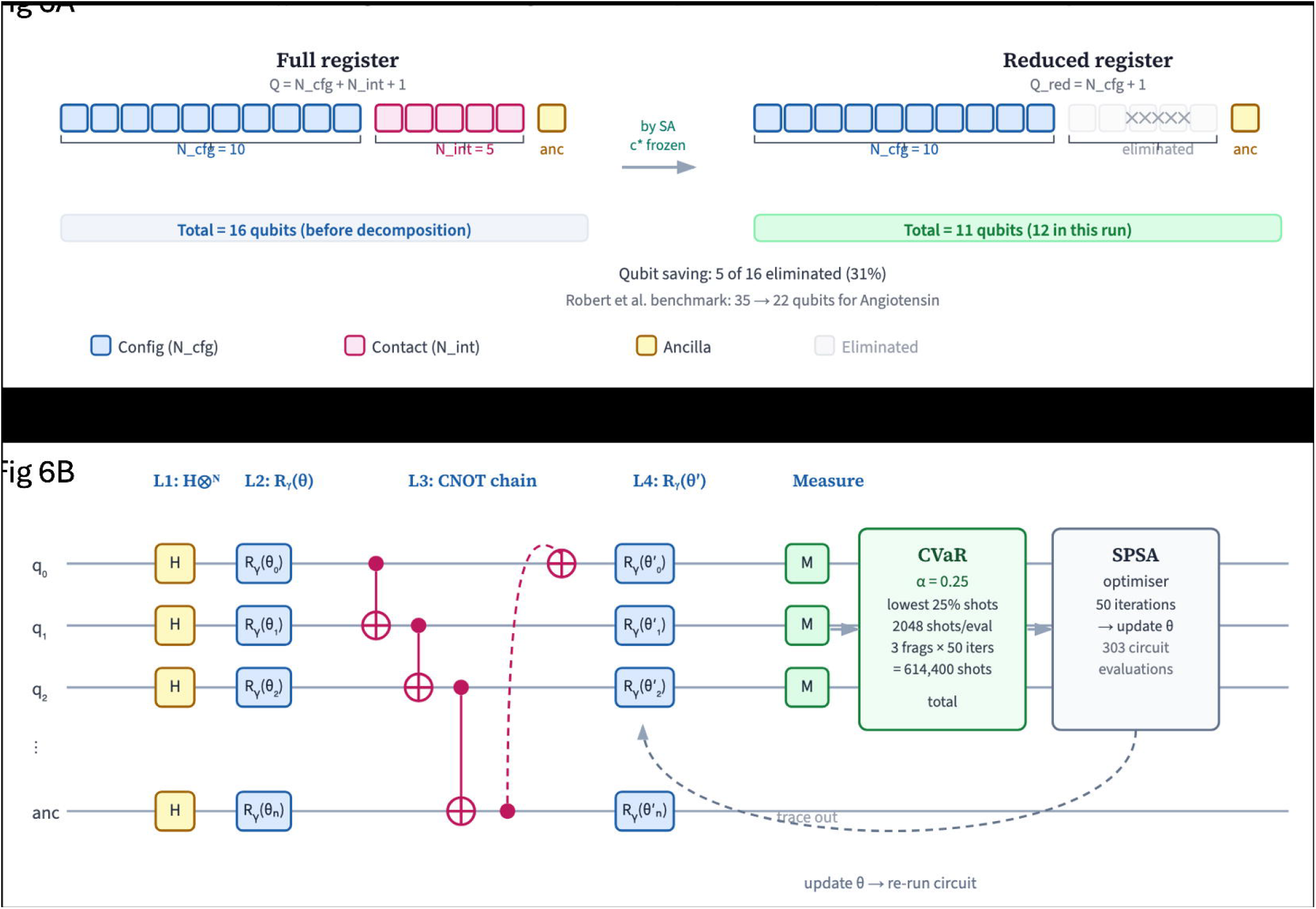
Qubit register and VQE circuit. A) Full (Q = 16) vs. reduced (Q_red_ = 11) register after freezing c* with SA. 12 qubits used. B) Hardware-efficient VQE ansatz circuit architecture (depth = 2).

### 4.3 The CVaR objective and NISQ-era robustness

QuPepFold uses the Conditional Value-at-Risk (CVaR) objective rather than the plain expectation value, consistent with recent variational literature. The CVaR aggregator was introduced by Barkoutsos, Nannicini and colleagues, who showed that optimising over the lowest-α tail of the measured energy distribution accelerates convergence and improves the probability of recovering the true ground state in combinatorial problems, precisely because the optimiser is rewarded for producing the best observed bitstrings rather than a high-variance mean [24]. Our benchmarking is consistent with that body of work: the CVaR-optimised architecture reached ground-state conformations roughly 30% faster than an expectation-value baseline, and reproduced absolute ground-state energies with greater than 90% fidelity on IonQ Aria trapped-ion hardware [12]. The CVaR principle has since been carried into domains well beyond chemistry, including variational optimisation for vehicular networks [29], confirming that CVaR is a validated, broadly applicable component of the variational toolkit. The pairing of CVaR with forbidden-state filtering—assigning infinite energy to self-intersecting walks— further sharpens the effective landscape that the optimiser sees, an approach broadly comparable to constraint penalisation in lattice-annealing formulations.

### 4.4 Energetics, contact potentials, and the scoring-function gap

The choice of the Miyazawa–Jernigan (MJ) statistical contact potential to score residue– residue interactions places QuPepFold in continuity with essentially all coarse-grained lattice approaches, classical and quantum alike [25,26]. The MJ matrices encode the dominant hydrophobic burial signal economically, which is what makes them tractable on a qubit register. The central finding of this study—that a favourable internal energy (E = −3.70) coexists with a poor 4.65 Å global RMSD—is best understood as an instance of the well-documented scoring-function gap that pervades the coarse-grained literature. A knowledge-based contact potential is a smoothed, low-resolution proxy for the true potential of mean force; its global minimum need not coincide with the experimentally observed backbone, particularly for a peptide such as the p53 fragment whose 1YCR structure is stabilised by binding to MDM2 and by crystal packing rather than by intrinsic free-solution folding. The discrepancy we observe is therefore not unique to the quantum solver: it is a property of the energy model, and the same limitation has been repeatedly noted in classical HP- and MJ-based lattice folding. Put differently, the quantum optimiser is doing its job correctly; the bottleneck is the accuracy of the Hamiltonian, not the solver.

This interpretation is corroborated by the substructure analysis. The improvement from a 4.65 Å full-backbone RMSD to 2.90 Å over a trimmed 10-residue subset indicates that segments of the chain are recovered with moderate local accuracy while the proline-containing central DLWKLLP fragment is poorly resolved. Because proline imposes severe φ-angle restriction that the four-state discrete turn alphabet approximates only crudely, the fragment assembly stage inherits incompatible local solutions—a failure mode of dynamic-programming stitching that is consistent with reports from fragment-based classical assembly as well. The locality and distance penalties (λ_dis, λ_loc), which earlier formulations set aggressively to 720.0 and 20.0 and which the present package disables by default to let MJ energies dominate, sit precisely at this tension between geometric feasibility and contact-driven folding.

### 4.5 Hardware validation in context

The successful execution on IBM_fez with greater than 90% energy fidelity is a meaningful demonstration, and it is worth contextualising. Trapped-ion platforms offer all-to-all connectivity and long coherence times that are well matched to the dense entangling structure of the hardware-efficient ansatz used here shown in (Fig 6B), in contrast to the limited graph connectivity of superconducting annealers that forces minor embedding. That QuPepFold produced faithful ground-state energies on real hardware—rather than only on statevector simulation—places it among the relatively small set of folding studies validated beyond classical emulation, alongside the Angiotensin VQE work [12] and the D-Wave annealing demonstrations [8]. At the same time, the gap between energy fidelity on hardware and structural fidelity against an experimental reference re-emphasises that hardware success and biological accuracy are distinct axes of evaluation.

### 4.6 Limitations, scope, and future directions

Several limitations frame the contribution and align with directions emerging across the field. First, the discrete four-state turn alphabet and 2D square-lattice tracing used during search are deliberately coarse; while they make the problem qubit-affordable, they cannot capture the continuous backbone geometry that distinguishes near-native from approximate folds, and the NeRF-like reconstruction to 3D cannot recover detail that the lattice never encoded. Second, the reliance on a single static contact topology from the SA pass, although it yields the decisive qubit saving, forecloses the conformational-ensemble view that is arguably the most compelling reason to study IDRs in the first place. Third, the scoring-function gap discussed above means that energy ranking alone is an unreliable proxy for structural correctness on this class of peptide.

These limitations point to concrete and literature-aligned next steps. Evaluating ensembles of low-energy conformers rather than a single final structure would align the output with the ensemble character of disordered peptides and with the way classical replica-exchange and Markov-state analyses report results. Enriching the turn alphabet or moving to finer lattices— at a controlled qubit cost—would narrow the geometric approximation gap, while problem-specific or symmetry-aware ansatz design could improve the VQE refinement of difficult proline-containing fragments. Most importantly, coupling the quantum search to a higher-resolution or learned scoring function, rather than the bare MJ potential, would directly attack the dominant source of the observed structural error. Together, these steps would advance hybrid quantum–classical folding from computational proof-of-concept toward practical utility for the intrinsically disordered targets that data-driven predictors currently handle poorly [4,5,11].

QuPepFold [10] draws on the methodological consensus in quantum-folding: resource-efficient register compression following Robert et al. [12], hybrid decomposition as recommended by Babbush and co-workers, CVaR-based optimisation introduced by Barkoutsos et al. [27], and MJ statistical scoring shared with the annealing tradition of Perdomo-Ortiz et al. [28]. The result is a fully modular pipeline validated on trapped-ion hardware. The 4.65 Å RMSD is consistent with the current state of the field: quantum-enabled folding works computationally and runs on real hardware, but structural accuracy is limited by the energy model, not the quantum solver.

## Acknowledgement

We acknowledge the use of IBM Quantum Credits for this work. The views expressed are those of the authors, and do not reflect the official policy or position of IBM or the IBM Quantum team. The authors would like to acknowledge the support and insights from Dr. Arindam Sarkar from the Department of Computing, MIT Vishwaprayag University. A heartfelt thanks to Suman A for the unconditional support. A warm heartfelt thanks to the staff and administration at the R V College of Engineering for their support.

## Code and Data availability

The code is available on GitHub and can be accessed via https://github.com/akshayuttarkarc/QuPepFold/tree/QA-VQE and data used for analysis is provided in the manuscript.

## 7. Supplementary information

Supplementary figure 1: QUBO degree-reduction: auxiliary variable substitution for quintic terms.

Supplementary table 1: DP assembly table for three fragments in strict mode. Blue rows = selected candidates. Post-stitch refinement: +8.73 → −2.58 MJ. Global SA (−3.703) selected as final output.

## References

1. Finkelstein, A. V., Bogatyreva, N. S., Ivankov, D. N., & Garbuzynskiy, S. O. (2022). Protein folding problem: enigma, paradox, solution. Biophysical Reviews, 14(6), 1255–1272. doi:10.1007/s12551-022-01000-1

2. Ivankov, D. N., & Finkelstein, A. V. (2020). Solution of Levinthal’s Paradox and a Physical Theory of Protein Folding Times. Biomolecules, 10(2), 250. doi:10.3390/biom10020250

3. Hayes, B. (1998). Prototeins. American Scientist, 86(3), 216. doi:10.1511/1998.25.216

4. Thilagavathi, N., & Amudha, T. (2014, March). An Analytical Study of NP-Hard Protein Folding Problems. 2014 International Conference on Intelligent Computing Applications, 184–188. doi:10.1109/icica.2014.47

5. A Perspective on Protein Structure Prediction Using Quantum Computers (n.d.-b). doi:10.1021/acs.jctc.4c00067.s001

6. Mustafa, H., Morapakula, S. N., Jain, P., & Ganguly, S. (2022, October 13). Variational Quantum Algorithms for Chemical Simulation and Drug Discovery. 2022 International Conference on Trends in Quantum Computing and Emerging Business Technologies (TQCEBT), 1–8. doi:10.1109/tqcebt54229.2022.10041453

7. Uttarkar, A., & Niranjan, V. (2024). Quantum synergy in peptide folding: A comparative study of CVaR-variational quantum eigensolver and moleculardynamics simulation. International Journal of Biological Macromolecules, 273, 133033. doi:10.1016/j.ijbiomac.2024.133033

8. Atari, M., & Majd, N. (2022, February 23). 2D HP protein folding using quantum genetic algorithm. 2022 27th International Computer Conference, Computer Society of Iran (CSICC), 1–8. doi:10.1109/csicc55295.2022.9780478

9. Irbäck, A., Knuthson, L., Mohanty, S., & Peterson, C. (2022). Folding lattice proteins with quantum annealing. Physical Review Research, 4(4).doi:10.1103/physrevresearch.4.043013

10. Uttarkar A, Niranjan V, Saxena A, Kumar V (2026) QuPepFold: A python package for hybrid quantum-classical protein folding simulations with CVaR-optimized VQE. PLoS One 21(2): e0342012. 10.1371/journal.pone.0342012

11. Robert, A., Barkoutsos, P. K., Woerner, S., & Tavernelli, I. (2021). Resource-efficient quantum algorithm for protein folding. Npj Quantum Information, 7(1).doi:10.1038/s41534-021-00368-4

12. A. Uttarkar, A. S. Setlur and V. Niranjan, “T-Gate Enabled Fault-Tolerant Ansatz Circuit Design for Variational Quantum Algorithms in Peptide Folding on Aria-1,” 2024 Global AI Summit, pp. 1271–1276, doi: 10.1109/GlobalAISummit62156.2024.10947993. https://ieeexplore.ieee.org/document/10947993

13. Uttarkar, A., Niranjan, V. A comparative insight into peptide folding with quantum CVaR-VQE algorithm, MD simulations and structural alphabet analysis. Quantum Inf Process 23, 48 (2024). 10.1007/s11128-024-04261-9

14. Rohl CA, Strauss CE, Misura KM, Baker D. Protein structure prediction using Rosetta. Methods Enzymol. 2004;383:66–93. doi:10.1016/S0076-6879(04)83004-0

15. Baek M, DiMaio F, Anishchenko I, et al. Accurate prediction of protein structures and interactions using a three-track neural network. Science. 2021;373(6557):871–876. doi:10.1126/science.abj8754

16. Jumper, J., Evans, R., Pritzel, A. et al. Highly accurate protein structure prediction with AlphaFold. Nature 596, 583–589 (2021). 10.1038/s41586-021-03819-2

17. Perdomo-Ortiz, A., Dickson, N., Drew-Brook, M., Rose, G., & Aspuru-Guzik, A.(2012). Finding low-energy conformations of lattice protein models by quantum annealing. Scientific Reports, 2(1). doi:10.1038/srep00571

18. Bahar, I., & Jernigan, R. L. (1997). Inter-residue potentials in globular proteins and the dominance of highly specific hydrophilic interactions at close separation 1 1 Edited by B. Honig. Journal of Molecular Biology, 266(1), 195–214.doi:10.1006/jmbi.1996.0758

19. Atari, M., & Majd, N. (2022, February 23). 2D HP protein folding using quantum genetic algorithm. 2022 27th International Computer Conference, Computer Society of Iran (CSICC), 1–8. doi:10.1109/csicc55295.2022.9780478

20. Irbäck, A., Knuthson, L., Mohanty, S., & Peterson, C. (2022). Folding lattice proteins with quantum annealing. Physical Review Research, 4(4).doi:10.1103/physrevresearch.4.043013

21. Bornberg-Bauer, E. (1996). Simple folding model for HP lattice proteins. Lecture Notes in Computer Science, pp. 125–136. doi:10.1007/bfb0033211

22. Irbäck, A., Knuthson, L., Mohanty, S., & Peterson, C. (2024). Using quantum annealing to design lattice proteins. Physical Review Research, 6(1).doi:10.1103/physrevresearch.6.013162

23. Nambury, S. B. (2026, June 23). Quantum Chemical Calculations of H,HeH<sup>+</sup>, LiH, BeH, H O, NH3, CH4, CO,and N Molecules Using the Variational Quantum Eigensolver(VQE): Ground State Energies,Potential Energy Surfaces, and Quantum Circuit Analysis. doi:10.21203/rs.3.rs-9953338/v1

24. A. Uttarkar and V. Niranjan, “Quantum Enabled Protein Folding of Disordered Regions in Ubiquitin C Via Error Mitigated VQE Benchmarked on Tensor Network Simulator and Aria 1,” IEEE Transactions on Molecular, Biological, and Multi-Scale Communications, doi: 10.1109/TMBMC.2025.3600516. https://ieeexplore.ieee.org/document/11130538

25. Bahar, I., & Jernigan, R. L. (1997). Inter-residue potentials in globular proteins and the dominance of highly specific hydrophilic interactions at close separation 1 1 Edited by B. Honig. Journal of Molecular Biology, 266(1), 195–214.doi:10.1006/jmbi.1996.0758

26. Zeng, H., Liu, K.-S., & Zheng, W.-M. (2012). The Miyazawa-Jernigan Contact Energies Revisited. The Open Bioinformatics Journal, 6(1), 1–8.doi:10.2174/1875036201206010001

27. I. O. Sokolov, P. K. Barkoutsos, L. Moeller, P. Suchsland, G. Mazzola, and I. Tavernelli, Microcanonical and finite temperature ab initio molecular dynamics simulations on quantum computers, 2020, arXiv:2008.08144 [quant-ph].

28. Perdomo-Ortiz, A., O’Gorman, B., Fluegemann, J. et al. Determination and correction of persistent biases in quantum annealers. Sci Rep 6, 18628 (2016). 10.1038/srep18628

