## Supplementary material for "Scalable Decoupling Contact and Configuration Spaces in a Quantum-Enabled Protein Folding Workflow": Supplemetary information

Supplementary figure 1: QUBO degree-reduction: auxiliary variable substitution for quintic terms

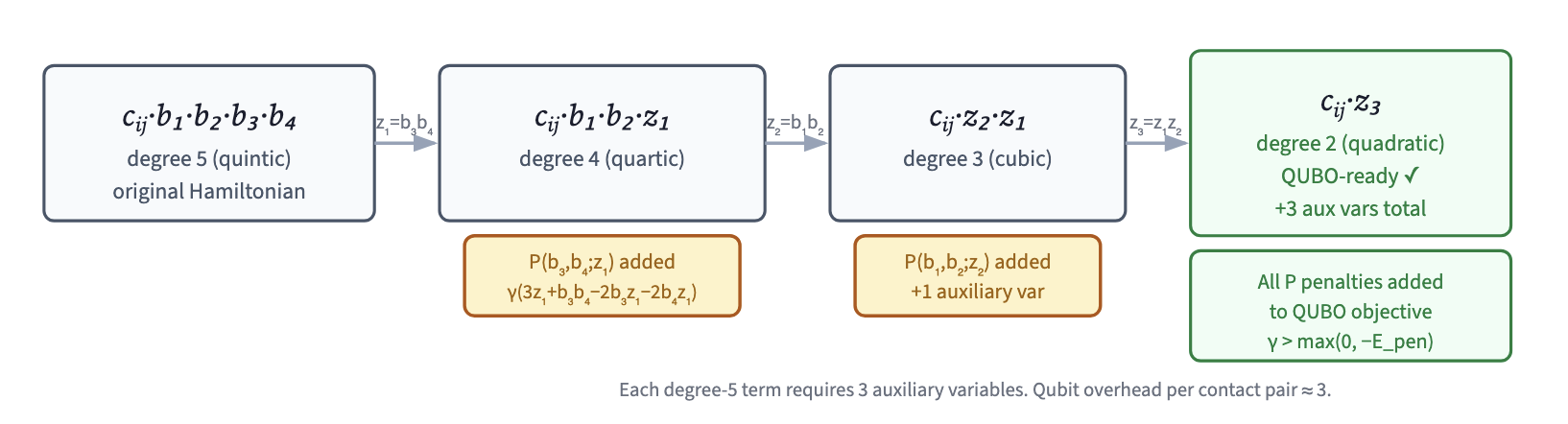

Supplementary table 1: Dynamic-programming fragment stitching trace

| **fragment_id** | **fragment_sequence** | **start_res** | **end_res** | **candidate_rank** | **bitstring** | **energy** | **probability** | **source** |
| --- | --- | --- | --- | --- | --- | --- | --- | --- |
| 1 | ETFSDLW | 0 | 7 | 1 | 100101000001 | -4.2057 | 0.070801 | vqe_sampling |
| 1 | ETFSDLW | 0 | 7 | 2 | 100100010001 | -3.462 | 1 | warm_start_sa |
| 1 | ETFSDLW | 0 | 7 | 3 | 110100010001 | -3.462 | 0.000244 | energy_table |
| 1 | ETFSDLW | 0 | 7 | 4 | 100110010001 | -3.462 | 0.000244 | energy_table |
| 1 | ETFSDLW | 0 | 7 | 5 | 110001100001 | -3.462 | 0.000244 | energy_table |
| 1 | ETFSDLW | 0 | 7 | 6 | 1000100001 | -3.462 | 0.000244 | energy_table |
| 1 | ETFSDLW | 0 | 7 | 7 | 110110010001 | -3.462 | 0.000244 | energy_table |
| 1 | ETFSDLW | 0 | 7 | 8 | 1000010001 | -3.462 | 0.000244 | energy_table |
| 1 | ETFSDLW | 0 | 7 | 9 | 110100100001 | -3.462 | 0.000244 | energy_table |
| 1 | ETFSDLW | 0 | 7 | 10 | 1001100001 | -3.462 | 0.000244 | energy_table |
| 2 | DLWKLLP | 4 | 11 | 1 | 100100001 | 0 | 1 | warm_start_sa |
| 2 | DLWKLLP | 4 | 11 | 2 | 100010001 | 0 | 0.000244 | energy_table |
| 2 | DLWKLLP | 4 | 11 | 3 | 110010001 | 0 | 0.000244 | energy_table |
| 2 | DLWKLLP | 4 | 11 | 4 | 101100001 | 1.2256 | 0.000244 | energy_table |
| 2 | DLWKLLP | 4 | 11 | 5 | 110100001 | 2.0059 | 0.000244 | energy_table |
| 2 | DLWKLLP | 4 | 11 | 6 | 110000001 | 5 | 0.000244 | energy_table |
| 2 | DLWKLLP | 4 | 11 | 7 | 101000001 | 9.4561 | 0.000244 | energy_table |
| 2 | DLWKLLP | 4 | 11 | 8 | 100000001 | 10 | 0.000244 | energy_table |
| 2 | DLWKLLP | 4 | 11 | 9 | 101111110 | inf | 0.533203 | vqe_sampling |
| 2 | DLWKLLP | 4 | 11 | 10 | 100100101011 | inf | 0.04834 | vqe_sampling |
| 3 | LLPEN | 8 | 13 | 1 | 11001 | -1.3519 | 1 | warm_start_sa |
| 3 | LLPEN | 8 | 13 | 2 | 10010 | -1.3519 | 0.003906 | energy_table |
| 3 | LLPEN | 8 | 13 | 3 | 11000 | -1.3519 | 0.003906 | energy_table |
| 3 | LLPEN | 8 | 13 | 4 | 10001 | -1.3519 | 0.003906 | energy_table |
| 3 | LLPEN | 8 | 13 | 5 | 10000 | 3.6481 | 0.003906 | energy_table |
| 3 | LLPEN | 8 | 13 | 6 | 10110 | 6.7949 | 0.003906 | energy_table |
| 3 | LLPEN | 8 | 13 | 7 | 10100 | 6.7949 | 0.003906 | energy_table |
| 3 | LLPEN | 8 | 13 | 8 | 11010 | 8.9094 | 0.003906 | energy_table |
| 3 | LLPEN | 8 | 13 | 9 | 11001011 | inf | 0.124756 | vqe_sampling |
| 3 | LLPEN | 8 | 13 | 10 | 10110100 | inf | 0.083496 | vqe_sampling |
